# Outer membrane vesicles from commensal and pathogenic anaerobic bacteria: a systematic review of literature reviews

**DOI:** 10.1101/2023.11.21.568143

**Authors:** Priscilla Castro-Vargas, Frédérique Barloy-Hubler, Luis Acuña-Amador

## Abstract

Outer membrane vesicles (OMVs) are constitutively produced by Gram-negative bacteria (GNB), those from pathogenic bacteria play key roles in pathogen-host interactions, modulating host’s immune response and promoting virulence. OMVs of commensal bacteria are fundamental in the maturation of the host immune system and to maintain intestinal homeostasis.

The objective of this systematic review was to synthesize the knowledge available on literature reviews on OMVs from anaerobic GNB. The information was classified into categories: induction and biogenesis, OMVs liberation, internalization by host cells, and their interaction. The most studied OMVs are from *Porphyromonas gingivalis* and *Bacteroides* spp.

## Introduction

Extracellular Vesicles (EVs) are a heterogeneous group of proteolipid bi-layered particles, 20-500 nm in size (1–7), secreted by most eukaryotic cells (fungi (8), protozoa (9), plants (10) and mammals (11)); archaea (12); and bacteria. EVs are released under both normal and pathological conditions by metabolically active cells (13).

Bacterial Extracellular Vesicles (BEVs) range between 20 and 500 nm in diameter, similarly to other EVs (13). BEVs were first identified in the Gram-negative bacterium (GNB), *Escherichia coli*, and later, found in the Gram-positive *Staphylococcus aureus* (14,15), and in mycobacteria (16). At first considered an artifact of lysis or cell wall turnover, their release is now recognized as a secretion system that can transfer an extremely diverse range of cargo and establish cell-to-cell communication (17,18).

To study BEVs, isolation and purification are done by low-speed centrifugation and supernatant filtration to remove cells and debris. Then, ultracentrifugation (100 000 to 200 000g for 1-4 h) and/or precipitation (using different concentrations of salts or polymers) are performed to separate BEVs from soluble components released during bacterial lysis. To increase purity and concentrate BEVs, other purification steps may be necessary; for example, by density-gradient centrifugation (in iodixanol, sucrose or dextran), size-exclusion chromatography (gel filtration), ultrafiltration (using a 50-100 kDa cutoff, cellulose or polyethersulfone membrane filters and applying pressure, vacuum or centrifugal force) or tangential flow filtration (7,13,15,19–22). Some steps in the isolation and purification protocol might affect the BEVs’ integrity and/or activity. Thus, quality control of isolates is key for research (22,23).

There is not a consensus about EVs nomenclature and the term itself is a generic one that encompasses several vesicles produced by cells (24,25). Based on their biotechnical production process, BEVs could have several names, including “bacterial double-layered membrane vesicles” (26) or “protoplast-derived nanovesicles” (27), to cite only a few.

Naturally occurring BEVs can be classified into at least three types: Cytoplasmic Membrane Vesicles (CMVs or MVs) produced by monoderm (Gram-positive) bacteria, Outer-Inner Membrane Vesicles (OIMVs) produced by diderm (Gram-negative) bacteria, and Outer Membrane Vesicles (OMVs) also produced by diderm bacteria (15,18,28,29). Another term “explosive outer-membrane vesicles” (EOMVs) has been used for OMVs produced by “explosive cell lysis” (15,22,30).

MVs (or CMVs) seem to be produced at lower quantities than the other BEVs and are implicated in stress resistance, biofilm formation and immunity (31). OIMVs are produced at very low quantities, representing for many Gram-Negative species around 1% of BEVs production. The possession of outer membranes, plasma membranes, and cytoplasmic content characterizes OIMVs. These BEVs seem to be involved in horizontal gene transfer (6,29,32). Finally, OMVs derive from the outer membrane (OM) (28).

As the study of BEVs is still challenging in terms of isolation, purification, and individual analysis (19), its classification and nomenclature is not congruent throughout literature (25,33). In this review, all BEVs derived from GNB are termed OMVs and their biological features will be discussed. A special emphasis will be provided on OMVs derived from anaerobic GNB. To do so, information derived from review articles about anaerobic GNB’s OMVs will be analyzed, and examples will be commented, to introduce the uninitiated reader to the subject, as a first educational approach to a topic in full expansion.

## Materials and methods

The recommendations of PRISMA (34) were followed for the writing of this systematic review:

### 1. Search

PubMed and ScienceDirect databases were searched from the oldest records to 10/31/2023. The search was performed using the following constructs: “Outer Membrane Vesicles”, “Extracellular Vesicles” or “Bacterial Membrane Vesicles” AND “Anaerobic bacteria”, “Bacteroides”, “Porphyromonas”.

### 2. Selection criteria

Inclusion criteria were: (1) literature reviews, (2) written in English, (3) published in a peer-reviewed journal, and (4) full-text available. The exclusion criteria were: (1) it did not contain relevant information or (2) the OMVs were not a central topic in the article. Results were downloaded and duplicate records were removed.

### 3. Data extraction

The information was organized in summary tables (Table S1) that included the metadata of each bibliographic review and the relevant information for the present study. Each article was evaluated independently.

### 4. Data analysis

The information obtained from each study was organized into five categories: (1) induction of vesiculation and biogenesis, (2) release, (3) content, (4) internalization of OMVs by a host cell, and (5) interaction of the OMV with the host cell. Based on this information, a synthesis of knowledge was written in each of the sections described above (Figure 1).

**Figure 1.**
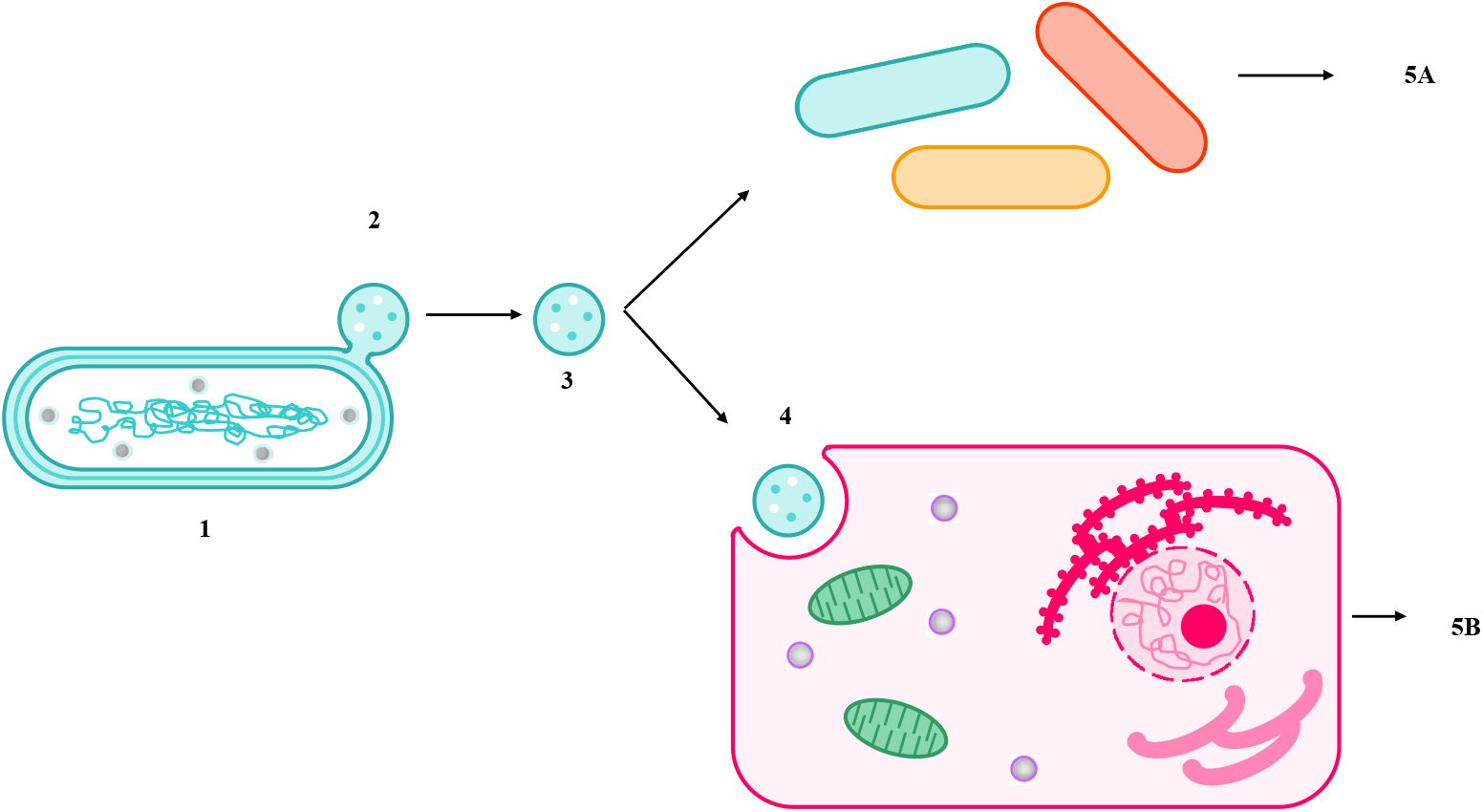
Summary of the paper’s sections. (1) Induction of vesiculation and biogenesis of OMVs. (2) Release of OMVs. (3) OMVs content. (4) Trafficking and internalization of OMVs. (5) Functions: (5.A.) Bacteria-bacteria interactions and (5.B.) Bacteria-eukaryotic interactions.

## Results

The search returned 1400 articles for inclusion in this review. After the process of selection, deduplication, and a manual review of each of the retrieved articles, a total of 53 review articles were used for this paper (Figure 2). The genera of Gram-negative anaerobic bacteria (GNB) whose OMVs have been studied in these articles correspond to *Porphyromonas* spp., *Bacteroides* spp., *Akkermansia* spp., *Aggregatibacter* spp., *Treponema* spp., *Fusobacterium* spp., *Tannerella* spp., *Odoribacter* spp. and *Prevotella* spp., in decreasing order of literature sources. None of the articles found had information on the OMVs of any other anaerobic GNB (Table 1). Notably, the only case for which an anaerobic GNB OMV was the primary topic of a review was for *Porphyromonas gingivalis* with 4 articles out of the 53 (7,5%) used in this review. The information collected from the articles was categorized into the following topics: (1) vesiculation induction and biogenesis, (2) release, (3) content, (4) trafficking and internalization of OMVs by host cells, and (5) interaction of OMVs with host cells. The information collected is detailed below.

**Figure 2.**
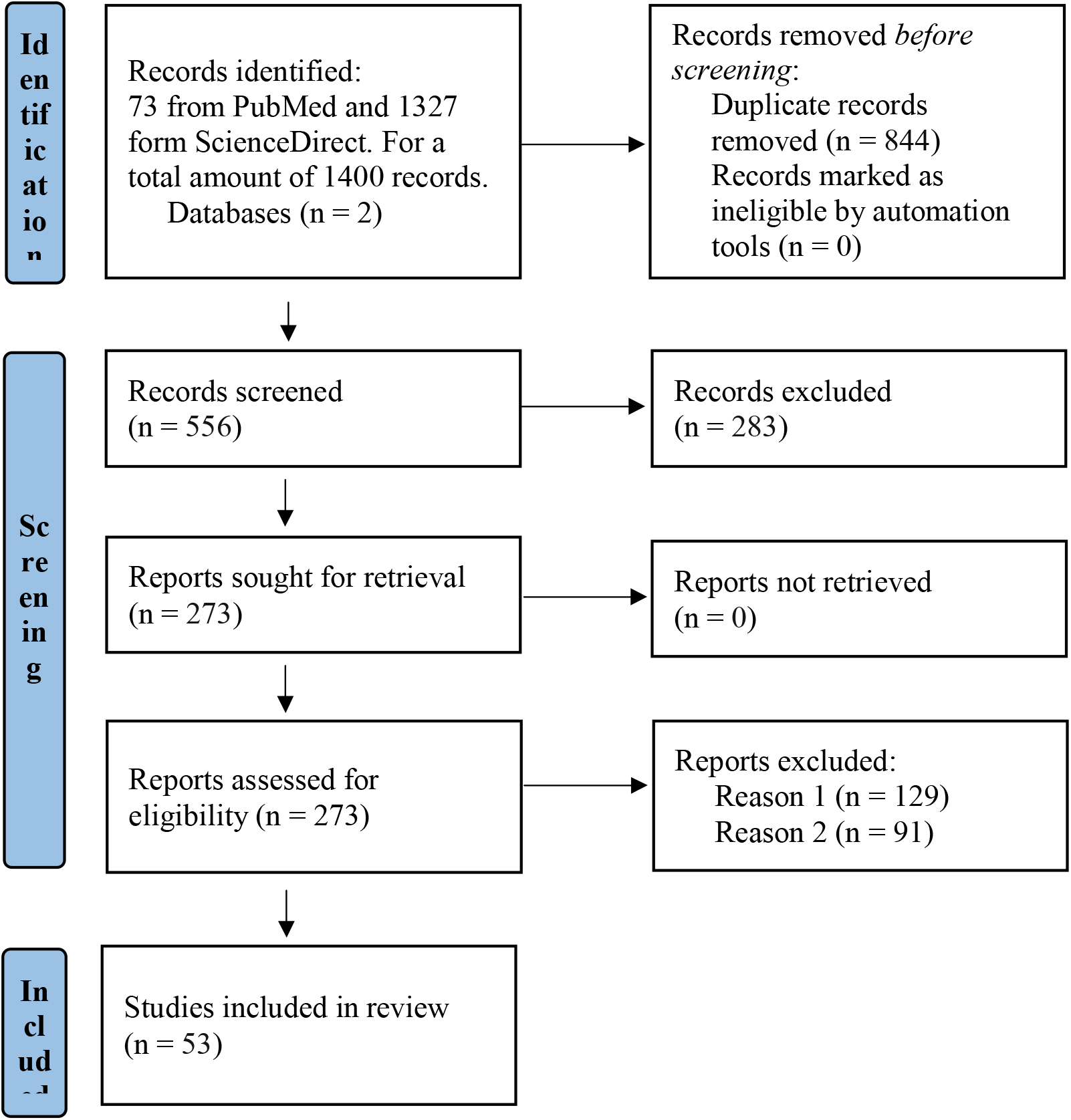
Flow chart based on PRISMA. Information on the article selection process for the systematic review is presented, along with the reasons for exclusion. Reason 1 for exclusion was that the article did not contain relevant information and Reason 2 was that OMVs were not a central theme in the article.

**Table 1.** Articles included in this review (number and percentage) with information on anaerobic GNB, presented by genus and with the reference number used throughout the main text of the review.

| **Bacterium** | **Article number** | **Article percentage** | **Reference in main review text** |
| --- | --- | --- | --- |
| *Porphyromonas* spp. | 36 | 67,9% | (15,18,41–44,46–51,22,53,54,56,58,59,61,64,66–68,29,70–73,78,79,33,35,36,38–40) |
| *Bacteroides* spp. | 33 | 62,3% | (6,7,47,50,51,55–57,59–61,63,15,64–66,68,71,73,74,77,78,81,18,82–84,29,37,43–46) |
| *Akkermansia* spp. | 16 | 30,2% | (6,7,64,72,78,79,81,83,15,18,22,39–41,44,50) |
| *Aggregatibacter* spp. | 13 | 24,5% | (15,22,79,81,82,37,51,55,56,62,64,70,78) |
| *Treponema* spp. | 7 | 13,2% | (39,40,44,58,59,64,66) |
| *Fusobacterium* spp. | 5 | 9,5% | (6,40,43,52,56) |
| *Tannerella* spp. | 2 | 3,8% | (40,59) |
| *Odoribacter* spp. | 2 | 3,8% | (18,81) |
| *Prevotella* spp. | 2 | 3,8% | (56,81) |
| Other anaerobic GNB | 0 | 0% | - |

### 1. Induction of vesiculation and OMV biogenesis

In GNB, vesicles are formed when a section of the outer membrane (OM) selectively buds, producing vesicles that encapsulate periplasmic material and detach from the OM. This results in the creation of spheroidal particles that consist of a single lipid bilayer (an outer layer rich in lipopolysaccharide (LPS) and an inner layer with more phospholipids) with a lumen (35–40).

OMVs share similar characteristics and biomolecular components to the precursor cells from which they originated (33,41,42). They range in size from 20 to 250 nm (42,43) and their composition reflects components of the OM and periplasm (33,41).

OMVs are produced constitutively, at low concentrations, during different growth phases and under different conditions such as in pure culture (solid or liquid media) or in natural environments, whether in biofilm or not (15,40,42,44–46). The rate of OMV production varies between bacterial species, even between strains of the same species, and can also be influenced by different environmental conditions (7,36,47). For example, among “the red complex”, a microbial group that colonizes periodontal pockets, *P. gingivalis* produces the most OMVs, followed by *Tannerella forsythia* and *Treponema denticola* (40). As mentioned, vesicle production also varies considerably between strains, for example in *Bacteroides fragilis*: whereas the OMVs were formed in great volumes by some clinical isolates, they were virtually completely absent in others (47).

Some factors such as quorum sensing, population size, and hostile or stress-generating external factors affect the properties of OMVs and increase their production (38,41,43,48–51). For instance, under low iron/heme conditions, *P. gingivalis* OMVs exhibited high levels of the protein HmuY, involved in heme acquisition (38). It is noteworthy that the molecular composition of naturally produced OMVs could differ from those induced by stress (42), and that OMVs can be isolated directly from the environment, particularly from aquatic environments, and have been isolated from household dust (46).

Despite initially being considered the result of cell lysis, vesiculation occurs in viable and metabolically active cells in which the integrity of the OM is not compromised (35,48,49). Moreover, certain OM or periplasmic proteins are more abundant in OMVs while others are completely absent (41). These observations support that vesiculation is regulated and that it would depend on environmental factors, bacterial pathogenicity, and the cellular metabolic state (36). However, experimental data to support the hypothesis that OMV biogenesis is a regulated process is lacking. Some important biological mechanisms that continue to be unknown are: genes and regulatory pathways involved in OMV production, and whether or not bacterial populations coordinate OMV liberation, to cite a few open questions.

Yet, OMV biogenesis is an energy-demanding process, so it should be a tightly controlled process, well-regulated and coordinated, as it enables crucial functions for the benefit of the producer (50). Some genes increase vesiculation rates by compromising bacterial OM, such as mutations in the Tol-Pal system or in *ompA* (43,49,52); for example, in *P. gingivalis* an *ompA* mutant showed hypervesiculation (53).

### 2. OMV release

Given the OMVs heterogeneity in production rate and size, it is likely that multiple biogenesis pathways occur in different stains and even in the same bacterial cell (50). It was observed that different *P. gingivalis* strains had varied vesiculation levels, correlating to fimbrial (FimA/FimR), OmpA-like protein and GalE expression (53,54).

OM microdomains formed during envelope biogenesis would exhibit compositional differences resulting in there being certain areas prone to form OMVs (36). The sites where OMV biogenesis would be favored would correspond to microdomains with specific physical characteristics in terms of curvature, charge, fluidity, and/or affinity (36).

Induction of OMV formation results from increased OM curvature, where OM and peptidoglycan (PG)-binding proteins are decreased, absent or broken (36,42,46,53,54). This mechanism may be concomitant with others that increase turgor pressure and promote OM curvature. For example, OMV production could be induced by a negatively charged molecule that induces repulsion of the OM anionic charge (46,55,56) and/or the physical force induced by accumulation of misfolded or overexpressed envelope proteins (46,53).

OM curvature is also favored in regions of low fluidity, where the proportion of saturated fatty acids is higher, which is also consistent with higher proportion of fatty acids in OMV envelopes (46). Additionally, the local curvature of bacterial OM is enhanced by extracellular signals (53), related to the affinity for amphipathic substances in the extracellular medium. This mechanism may be relevant to stress-induced responses (46).

Another factor that could promote OMV biogenesis is OM curvature increase, favored by conformation of transmembrane proteins (36,46). Finally, decreased, or absent expression of *vacJ* and/or *yrb* genes, due to iron and sulfate depletion, leads to the accumulation of phospholipids in OM which induces a change in membrane curvature (43,54,57).

As a consequence of the increase in OM curvature, by the mechanisms outlined above, a budding arises and then the OM constricts forming a proteo-liposome and encapsulating contents in its lumen (36,41,44). Different inductors of OMV production and release should influence different biogenesis pathways and induce mechanisms for OMV liberation from its parental cell. Knowledge about OMV surface biomarkers and their differential expression is needed. Without this information, the study of biogenesis routes or the mechanism of release would be extremely difficult.

### 3. OMVs content

OMVs composition varies even within the same bacterial population and depends on the growth phase and environmental conditions (48,52). However, detailed molecular analysis of OMV contents supports the idea of selective enrichment or exclusion of specific proteins, lipids and other biomolecules (46,49,58). Load selection mechanisms in OMVs remain incompletely understood (38,58). A schematic representation of the content of the OMVs is presented in Figure 3.

**Figure 3.**
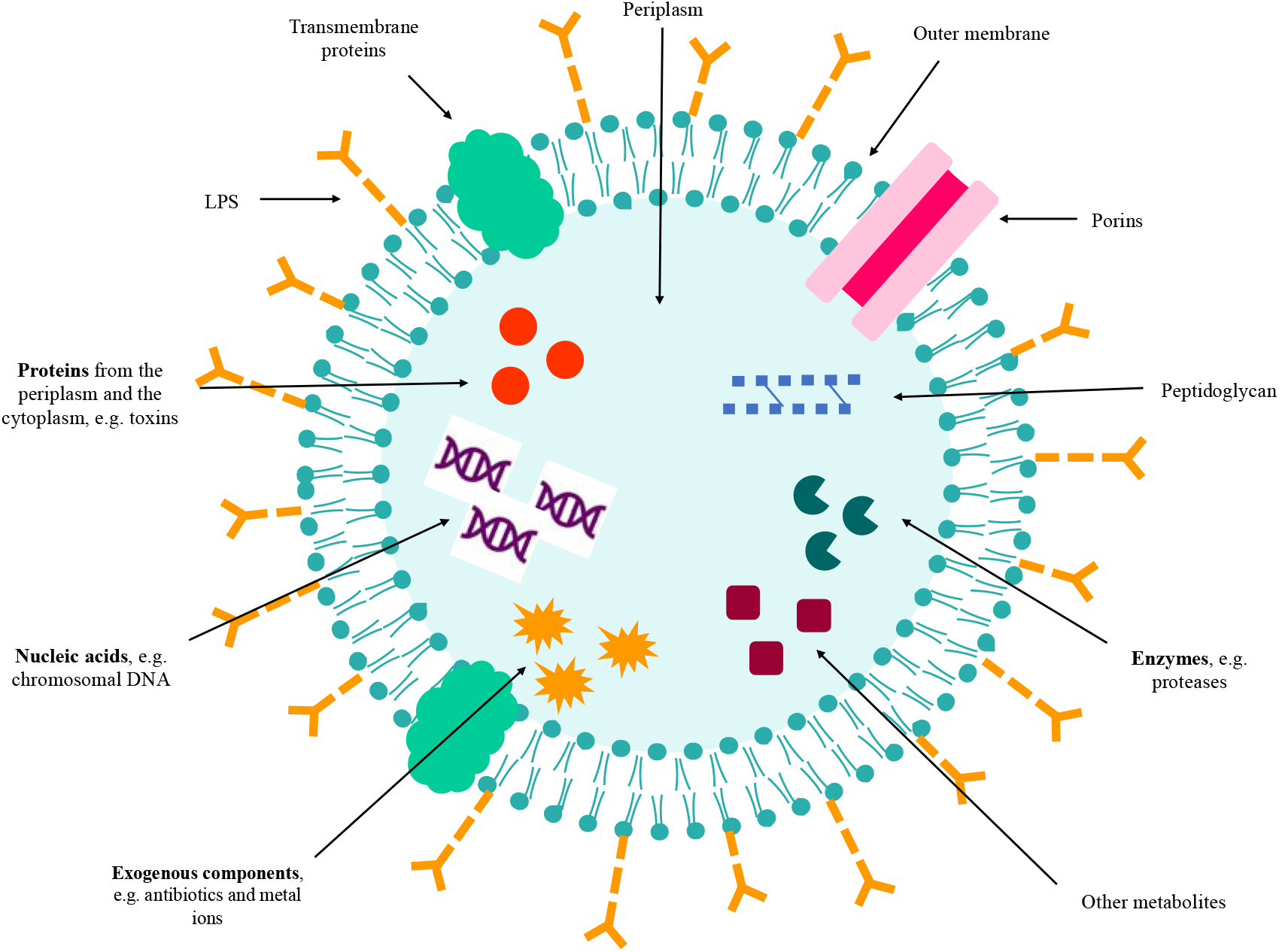
OMVs content and structure. OMVs are enveloped by an outer membrane bilayer of Gram-negative bacteria (GNB). They contain LPS, porins, transmembrane proteins, peptidoglycan, enzymes, proteins, nucleic acids, exogenous components, and other metabolites.

In the intravesicular lumen, the content or cargo is mostly composed of OM and periplasmic proteins, and to a minor degree, inner membrane (IM) and cytoplasmic proteins, including soluble proteins, integral membrane proteins, and lipoproteins (36,38,40,41,51,54,59,60). For instance, *Aggregatibacter actinomycetemcomitans* exports leukotoxin, a Type I-secreted, highly conserved repeat toxin (RTX) family-member, through OMVs (36), while TonB-dependent receptors have been found in vesicles of *T. forsythia* (40) and Type V secretion system proteins have been identified in OMVs from *Fusobacterium nucleatum* (40). It has also been described, in the lumen, genetic material including chromosomal DNA, rRNA, mRNA, miRNA lncRNA, circRNA and other small RNAs (35,39,48,54,55,58,61,62). For example, small RNAs have been found in vesicles from *A. actinomycetemcomitans, P. gingivalis* and *T. denticola* (39). In the case of *P. gingivalis,* chromosomal DNA, mRNA and rRNA are present in OMVs (48). Some of these biomolecules could be biologically active (58,63), and misfolded proteins or exogenous components such as antibiotics and metal ions have also been reported (48). OMVs from *B. fragilis* exhibited inhibitory activity against other Bacteroidales through the intravesicle antimicrobial peptide Bacteroidales secreted antimicrobial proteins (BSAP) (55).

Abundant structural proteins and porins, as well as enzymes, ion channels, transporters, and proteins related to stress responses are found in most OMVs in both the lumen and outer layers (55). For instance, several hydrolytic enzymes such as glycosidases, sulfatases and proteases are contained in vesicles derived from *Bacteroides* spp. (55). Moreover, OMV envelopes may contain a specific lipid profile that differs from OM, with distinctive differences in the LPS profile (46,50). *P. gingivalis* synthesizes anionic LPS and neutral LPS, both of which are present in OMVs; nonetheless, lipid A was deacylated in vesicles in comparison to the OM and cardiolipin was enriched in OMVs from *A. actinomycetemcomitans* (46,52). Likewise, OMVs contain pathogen-associated molecular patterns (PAMPs) present in the outer membranes of bacteria (49,64). For instance, polysaccharide derived from OMVs shed by *B. fragilis* is sensed by gut dendritic cells trough Toll-Like Receptor 2 (TLR2) and leads to Interleukin 10 (IL-10) expression (59). These interactions with host immune cells through PAMPs provide OMVs with immunomodulatory properties.

In this sense, virulence factors are associated with the OMVs released by pathogens (44,65). Thus, OMVs are a rich source of antigenic determinants and other bioactive components (66–68). Said biomolecules would play various roles in adherence and invasion to host cells, resistance to antibiotics, biofilm formation, promotion of virulence, or have immunomodulatory effects (7,29,36,63). Table 2 provides more details on the virulence factors, mainly proteases and adhesins, described in anaerobic GNB’s OMVs.

**Table 2.**
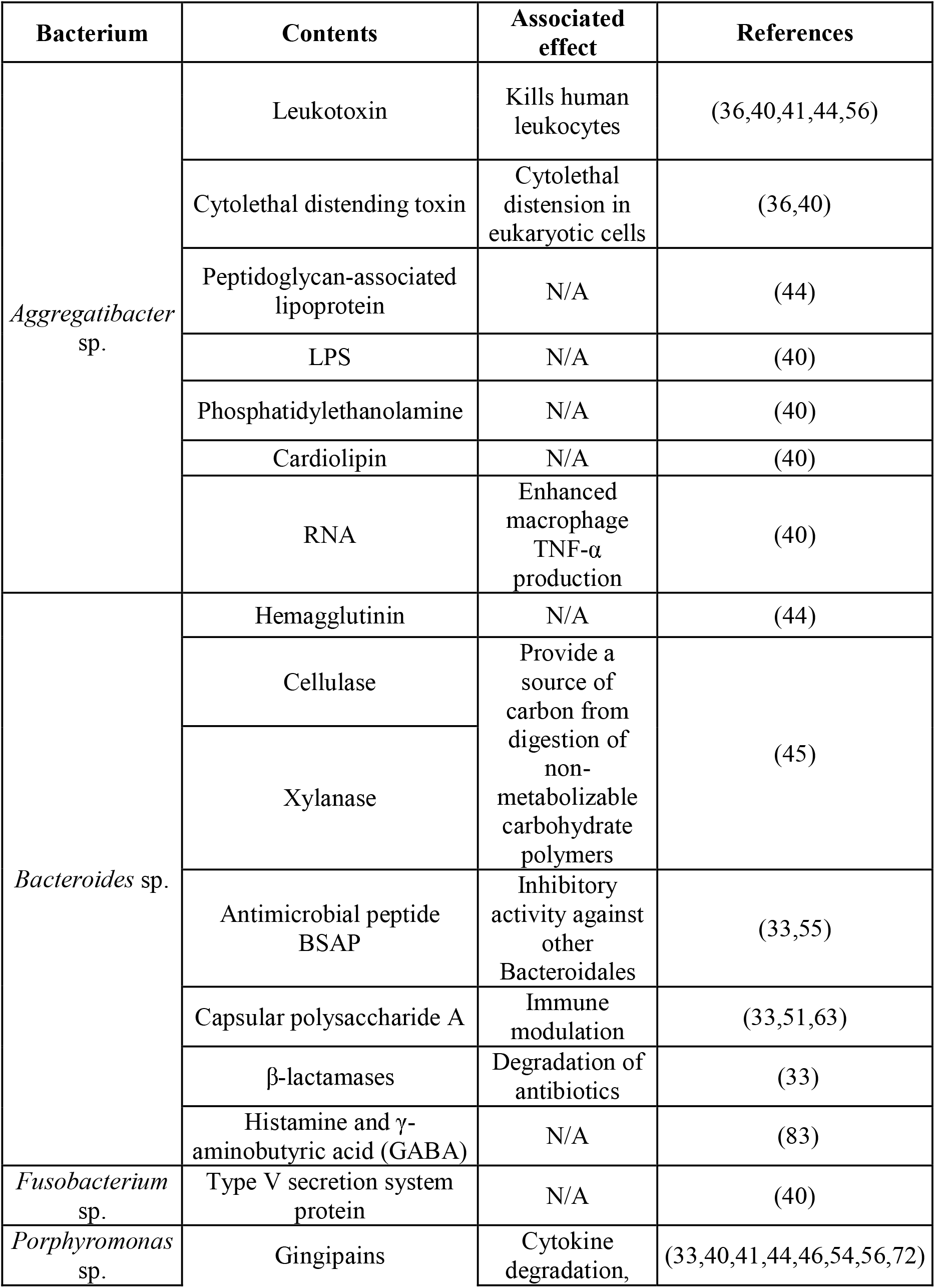
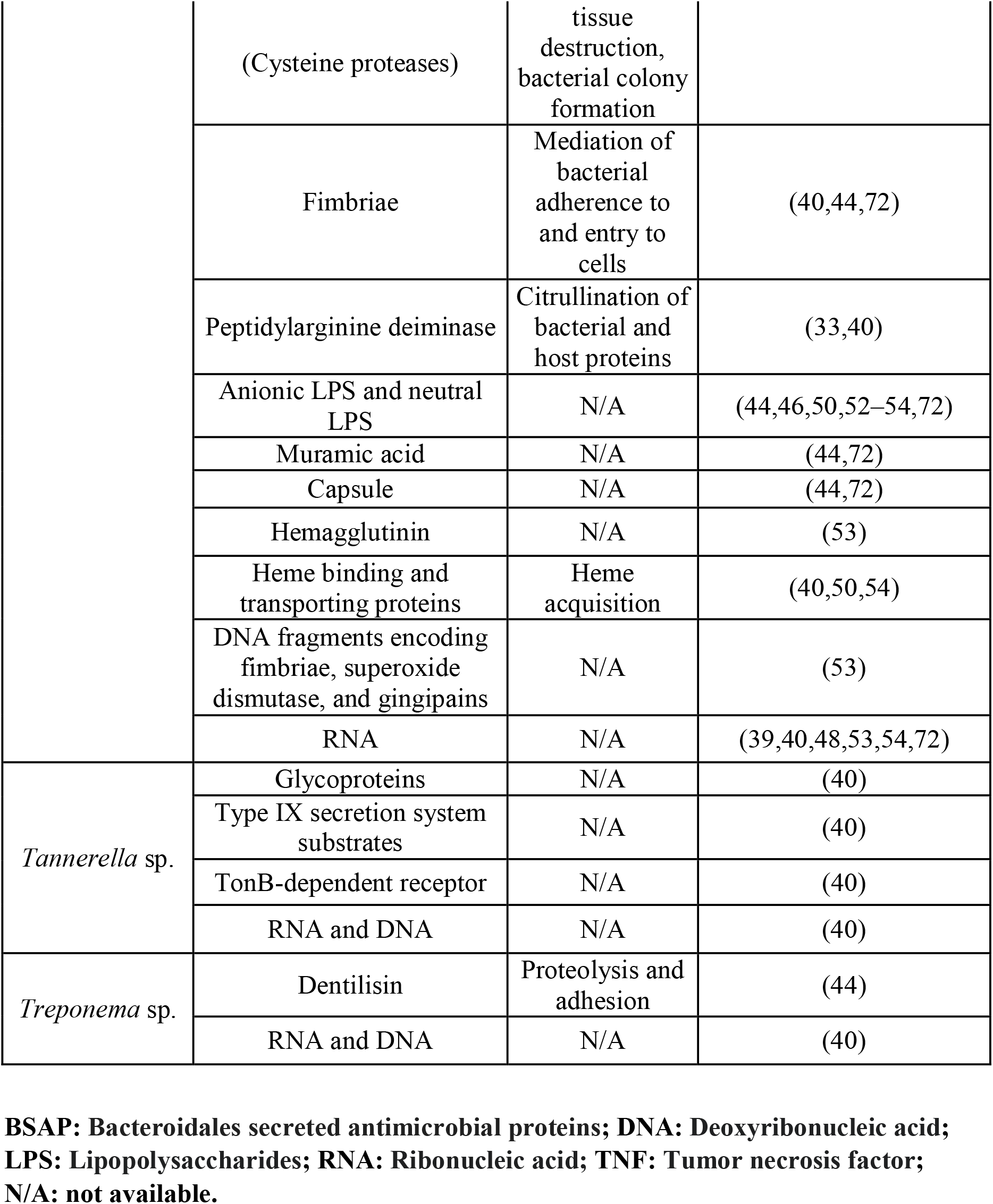
OMV-associated contents in anaerobic Gram-negative bacteria (GNB).

Other OMV-associated proteins may play roles in interspecies cooperation and intercellular communication (36). For example, OMVs from *Bacteroides* spp. contain proteins involved in carbohydrate (e.g. cellulase and xylanase) and amino acid metabolism, the citrate cycle, and cell division (45,60).

Many scientific teams, using proteomics, have extensively studied the presence of proteins from different cell compartments. However, protein sorting for cargo into the OMV lumen is poorly understood.

Additionally, the reported presence of nucleic acids in OMV’s lumen is more controversial since the biological mechanism is unknown and may be the result of contamination or a random event, rather than of active transport and packaging. It has been proposed that nucleic acids could be present in OMV’s lumen through the cytoplasmic route (where cytoplasmic content is trapped into OIMVs), through the periplasmic route (where nucleic acids are exported from the cytosol to the periplasm, and then packed into OMVs), and/or through a extracellular route (where OMVs reassemble in extracellular media and incorporate nucleic acids, and/or phages inject DNA into OMVs).

DNA and/or RNA sequencing from individual OMVs in a population might shed light on this controversy. Moreover, whether or not these nucleic acids are biologically significant and can contribute in the adaptation and response from host cells interacting with anaerobic GNB OMVs should be further explored.

### 4. Internalization of OMVs by a host cell

Once released, OMVs can travel to distant sites and interact with various cells; this has been studied mainly in pathogenic anaerobic GNB. OMVs lack a mechanism to self-target cells and the medium they are in would affect their transport (36,46). Several factors have been identified to promote OMVs adhesion to eukaryotic cells, such as LPS-binding proteins (46). In the case of *P. gingivalis,* fimbriae confer adhesive abilities to OMVs, and *T. denticola* OMVs also contain adhesins such as dentilisin (69).

A few routes for OMV entry into host cells have been proposed: (1) endocytosis, in eukaryotic cells it can be clathrin- or caveolin-dependent, (2) pinocytosis, an actin-induced process where nonspecific uptake of extracellular components occur, (3) by lipid rafts in the host plasma membrane via caveolins or GTPases, (4) phagocytosis, and (5) by direct membrane fusion (18,22,51,53,54,56,59,62–65,70,33,71–73,37–40,43,46,50). In that sense, OMVs have an intrinsic ability to fuse with the cell and spill their contents into the host’s cytoplasm (39,46,64,72,74).

Importantly, OMVs size and content have been shown to influence the route and speed of uptake, with potentially several endocytic pathways in simultaneous use (6,54,70,71,73). For example, rough OMVs with shorter LPS will rapidly fuse to the host membrane surface, whereas smooth OMVs, containing LPS with O-terminal antigen-terminal chains, tend to retain their spherical shape during interaction (71).

Importantly, for pathogens with an intracellular (facultative) lifestyle, OMVs can be released directly into the host cell without the need to overcome a cytoplasmic membrane. Therefore, host responses are likely to differ according to pathogen lifestyle and interaction pathway (39).

Additionally, OMVs from intestinal bacteria can be detected in host’s blood and urine. This suggests that OMVs can cross the intestinal epithelium and vascular endothelium to reach sites beyond the gastrointestinal tract (65). To do this, OMVs can use two distinct pathways: the paracellular or transcellular pathways (37,65). The paracellular pathway works trough the disruption of tight junctions (TJ) that form the intercellular barrier between epithelial cells; and the transcellular pathway consists in OMVs being internalized trough endocytosis, transported through the cell body and released at the basolateral cell surface (75,76). For example, OMVs produced by *Bacteroides thetaiotaomicron* have been reported to travel by both pathways, with imaging techniques showing that after hours of oral administration of OMVs, they can be detected in tissues such as the liver (76). These results support the possibility that OMVs can act as a long-distance communication system between microbiota and the host.

As for specific examples, it has been described that OMVs from *P. gingivalis* can be internalized via receptors, such as integrin α5β1, via lipid rafts and via caveolin/clathrin-independent endocytosis (40,50,53,54,64,71). OMVs from *P. gingivalis* have been shown to enter cells such as human oral keratinocytes and gingival fibroblasts more efficiently than parental bacterial cells (72). Moreover, the content of *T. denticola* OMVs can be sensed by the absent in melanoma 2 (AIM2) inflammasome and consequently trigger an inflammatory cell death (59).

Limited information about receptors for internalization and the actual entry route skew our understanding on anaerobic GNB OMVs entry into cells. Moreover, a bias exists in the lack of information for almost all anaerobic GNB (*P. gingivalis* being perhaps the exception), as they are not usually recognized as bacterial models of study, and therefore their particularities are poorly understood.

### 5. Interaction of OMVs with host cells

Host cells can internalize OMVs released by an anaerobic GNB. Host cells can be distant from where the OMVs parental cell is located. In animals, OMVs could be dispersed in body fluids (36,65,67). The host cells that internalize OMVs can be bacterial or eukaryotic, as detailed below.

#### 5.A. OMVs-bacteria interactions

OMVs play an important role in competitive and cooperative functions for the microorganisms that secrete them, and for the microbial community by participating in events such as: i. biofilm formation, ii. gene transfer, iii. antibiotic resistance, and iv. phage neutralization (35,36,38–40,42,55,70,71,77,78).

i. OMVs play their role in biofilm formation and stabilization by participating in nutrient acquisition, protection against antimicrobial effects, nucleation, and intercellular communication (35,41,70). It has even been reported that OMVs can be part of the biofilm matrix and are one of the most important protein sources for the matrix of certain biofilms (35,46,71). For example, under starvation conditions, quorum sensing and the HmuY protein detected in OMVs of *P. gingivalis* aid in cell survival and biofilm formation (36,70).
ii. OMVs could participate in the lateral transfer of plasmids and genomic DNA to other bacteria (35,37,46,55,70,71). Gene transfer via OMVs contributes to the propagation of factors necessary for bacterial adaptation, such as antimicrobial resistance genes or virulence factors (35,48,55,58,70). For example, chromosomal DNA transfer by OMVs from two different strains of *P. gingivalis* with subsequent integration by homologous recombination and gene expression in the recipient strain has been demonstrated (48).
iii. OMVs can protect bacteria from antibiotics in several ways: (1) by carrying β-lactamases (36,63,65,70,71), (2) by binding antibiotics in the extracellular environment, serving as a decoy, and preventing cellular entry (38,41,55,70,71), and (3) by exhibiting a high content of drug-binding proteins (70). In a mixed bacterial population, OMVs may provide antibiotic protection not only to parental cells, but to the whole bacterial population; thus, OMVs may be involved in the emergence of drug-resistant strains (36,70). For example, β-lactamases associated with OMVs from *Bacteroides* sp. provide antibiotic resistance to parental cells, but also to other commensals and pathogens against β-lactam antibiotics (55,63).
iv. Since OMVs have similar surface structures to the parental bacteria, the release of OMVs may act as a trap to prevent phage adsorption (70). Phages form complexes with OMVs and avoid binding to bacterial cells, preventing harmful effects on bacteria (55,70).

Other functions of OMVs have been described; for example, they could be a removal system for unnecessary or harmful substances that accumulate in microbial cells, such as misfolded proteins and xenobiotics (46,71). A graphical summary of these and other functions performed by OMVs is presented in Figure 4.

**Figure 4.**
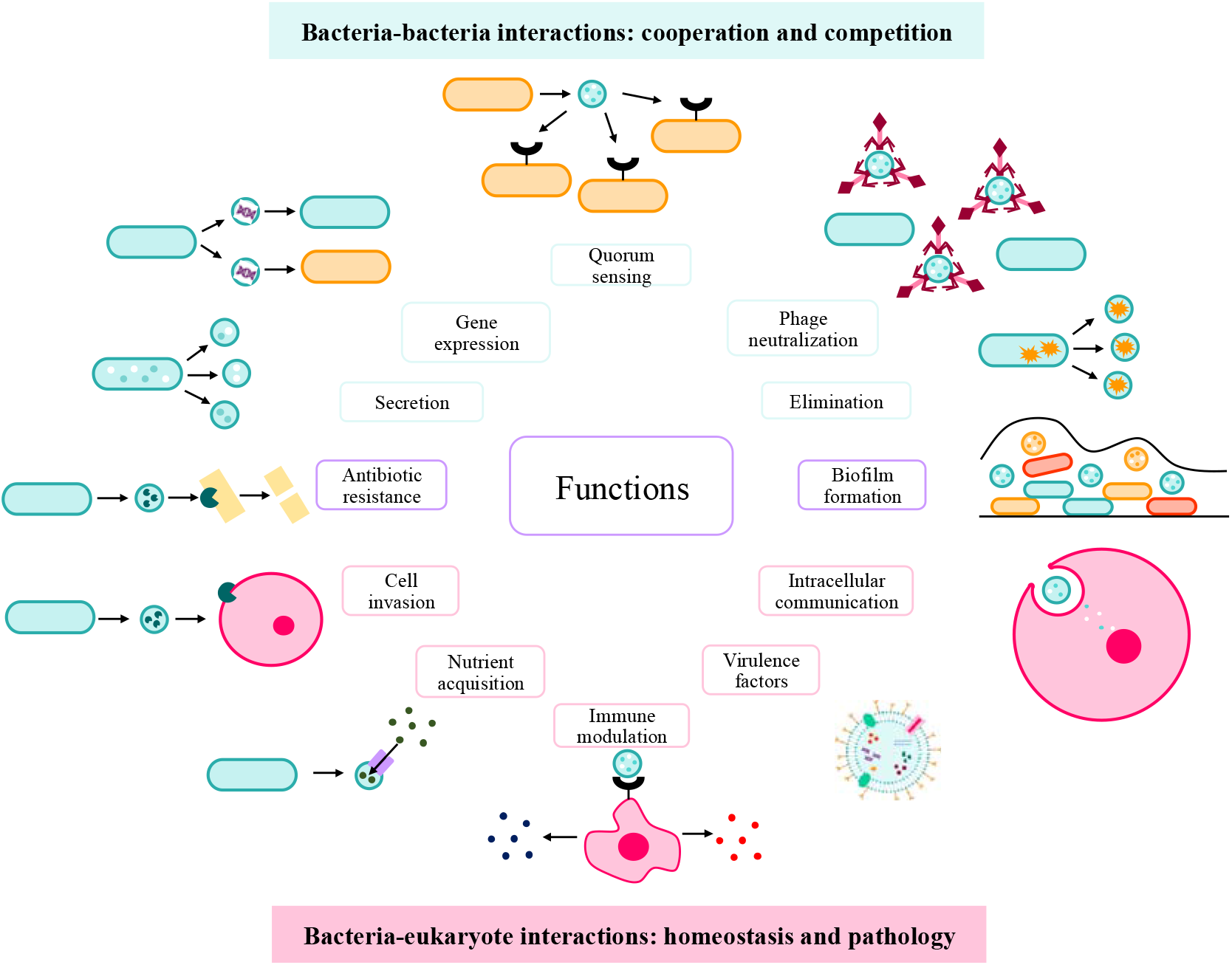
OMVs functions. OMVs play several roles in bacteria-bacteria interactions in processes of cooperation and competition, and in bacteria-eukaryotic interactions either in homeostasis or pathology.

#### 5. B. OMVs-eukaryotic cell interactions

OMVs of pathogens and commensals play a pivotal role in the communication between bacteria and their eukaryotic hosts (51,55,57,79). Generally, OMVs of pathogenic and commensal bacteria are related to distinct interactions with the host: i. OMVs of pathogenic bacteria participate in the pathological process, and ii. OMVs of commensal bacteria promote host homeostasis (47,55,79,80).

##### 5. B.1. OMVs from pathogenic bacteria-eukaryotic cell interactions

The content of OMVs of pathogenic bacteria is involved in host cell invasion, nutrient acquisition, antibiotic resistance, immune modulation, virulence, biofilm formation and intracellular communication (35,36,51,53–56,70,71). In addition to transferring toxins to host cells, OMVs contribute to pathogen survival and maintenance of pathogenesis by evading host immunity, aiding adaptation to the stressful environment, and nutrient capture (70). Some examples of virulence factors and their interaction with the host are cited in Table 2.

OMVs from different bacteria, and their components, induce distinct immune responses (78). LPS derived from OMVs activates toll-like receptor 4 (TLR4), which triggers inflammation through the nuclear factor kappa-light-chain-enhancer of activated B cells (NF-κB), proinflammatory cytokine production, caspase-11-dependent effector responses and cell death (35,59,64,70,73). Moreover, OMVs polysaccharides activate TLR2 and cytosolic nucleotide-binding oligomerization domain-containing protein 2 (NOD2), which induces phagocytosis and anti-inflammatory responses associated with autophagy (43,59). Finally, flagellin in OMVs is detected by the nucleotide-binding domain leucine-rich repeat (NLR) family, apoptosis inhibitory protein 5 (NAIP5) and activates the NLR family CARD domain-containing protein 4 (NLRC4) inflammasome, while OMV-derived dsDNA promotes the formation of the AIM2 inflammasome; both inflammasomes trigger caspase-1-mediated pyroptosis and inflammation (43).

An example is *P. gingivalis* OMVs which can induce inflammasome activation via caspase-1 activation and IL-1β and IL-8 production (54,59,72), and induce the production of tumor necrosis factor alpha (TNF-α), IL-12p70, IL-6, IL-10, interferon beta (IFNB) and nitric oxide in macrophages (54,72). In addition, gingipains in *P. gingivalis* OMVs induce IL-8 scission and activation, leading to neutrophil recruitment (78). Importantly, it has been reported that inflammation may be enhanced in macrophages exposed to *P. gingivalis* OMVs compared to inflammation induced by exposure to *P. gingivalis* cells (72).

The involvement of *P. gingivalis* OMVs in several systemic diseases has been described, for example in: (1) periodontitis: OMVs induce inflammation and tissue destruction, (2) diabetes: OMVs translocate to the liver and attenuate insulin sensitivity and glycogen synthesis, (3) arteriosclerosis: OMVs possess potent platelet aggregation activity, stimulate foam cell formation and low-density lipoprotein (LDL) aggregation, (4) rheumatoid arthritis: OMVs induce production of anti-citrullinated protein antibodies and loss of tolerance to citrullinated proteins, (5) Alzheimer’s disease: OMVs induce brain inflammation (22,40,54,72).

For other pathogens, the amount of information found is more limited. In the case of *T. denticola*, its OMVs could destroy the integrity of the epithelial cell barrier (64,66). *F. nucleatum* OMVs activate TLR2-MyD88-NF-kB signaling through the FomA porin, and thus exert proinflammatory effects and regulate innate immunity (43). Finally, *A. actinomycetemcomitans* OMVs contain cytolethal distending toxin and leukotoxin A, the latter of which is lethal to human monocytes and polymorphonuclear leukocytes, and the genetic material contained in OMVs could regulate human macrophage gene expression, promoting TNF-α production (40,44,81).

Limited information about OMVs cargo is presented in the literature that presents regulation of eukaryotic cells after OMVs content’s internalization. Besides the examples presented, and to add to the controversy about the presence of nucleic acids in OMVs, several studies presented downregulation of immune effectors after delivery of small RNAs into host cells by OMVs from *A. actinomycetemcomitans*, *P. gingivalis*, *T. denticola*, and/or *T. forsythia*. The lack of genetic tools to inhibit vesiculation in order to understand OMVs implication in infections is a difficulty that should be addressed.

##### 5. B.2. OMVs from commensal bacteria-eukaryotic cell interactions

OMVs from commensal bacteria can generate positive effects on host immune tolerance by regulating the host innate immune system, assisting in the maturation of the immune system, and promoting intestinal homeostasis (48,55,57,70,80,82). For example, OMVs would serve to control bacterial populations by loading with toxic molecules, such as peptidoglycan hydrolases, murein hydrolases, endopeptidases, and other types of enzymes that induce bacterial lysis, thus controlling bacterial overgrowth (45,55,57,71). In addition, OMVs from commensal intestinal bacteria promote regulatory T cells and anti-inflammatory cytokine secretion through activation of host dendritic cells (DCs) (57,66,73).

For example, *Akkermansia muciniphila* OMVs have been associated with intestinal barrier fortification and reduced inflammation by enhancing TJ expression and downregulating TLR expression in intestinal epithelial cells (6,37,64,81,83). In mice models, *A. muciniphila* OMVs have been described to improve TJ function and glucose tolerance (37,64,70,81,82), colitis symptoms and enhance intestinal epithelial barrier integrity (51,55,64), and provided protection against inflammatory bowel disease (IBD) phenotypes (70). Furthermore, direct intervention of mice with *A. muciniphila*-derived OMVs had exhibit an anti-osteoporotic effect (62,78). In addition, *Odoribacter splanchnicus*-derived OMVs have exhibited immunomodulatory properties, mitigating IL-8 production in enterocyte cultures treated with LPS from *E. coli* (18,81).

Moreover, polysaccharides associated with *B. fragilis* OMVs are detected by mouse intestinal DCs via TLR2 to induce IL-10 expression, which in turn enhances Treg function and decreases intestinal inflammation (48,51,55,59,65,81). Noticeably, *B. fragilis* OMVs have been reported to have a protective effect on some types of colitis (7,64); for example, they prevented the development of colitis in mice by inducing immune tolerance via DC activation (81). *B. fragilis*-derived OMVs are rich in polysaccharide A, which ameliorates human inflammatory disorders such as inflammatory bowel disease and multiple sclerosis. These OMVs show comparable therapeutic activity to their parental bacteria in terms of immune regulation (57,61).

*B. thetaiotaomicron* and *Phocaeicola vulgatus* (previously known as *Bacteroides vulgatus*) OMVs have also been reported to have immunomodulatory effects by controlling DC maturation, suppressing inflammatory cytokine secretion and promoting anti-inflammatory cytokine production (6,37,55,64). Specifically, *B. thetaiotaomicron*-derived OMVs contain enzymes involved in polysaccharide digestion (7,29,65,78) and mediate cholesterol uptake into enterocytes through positive regulation of transporters (55), but can also be taken up by intestinal macrophages and elicit an inflammatory response by inducing the production of proinflammatory cytokines and driving the development of colitis in susceptible mice (81).

The interaction between anaerobic GNB OMVs and eukaryotic cells is not fully known, many aspects need clarification, as stated throughout this review. In this sense, OMVs potentialities, especially for those produced by commensal anaerobic GNB, as drug-delivery systems or other biotechnological/biomedical uses are still too poorly understood for clinical applications.

## Concluding remarks and perspectives

OMVs are unique delivery systems used by GNB that fulfill several roles in bacteria-bacteria interactions and bacteria-eukaryotic interactions. This systematic review offers a synthesis of current knowledge available for the OMVs produced by anaerobic GNB. Induction of OMVs production, its biogenesis, contents, liberation from the parental cell, spread to the environment, internalization by either bacterial or eukaryotic host cells and the interactions with host cells are discussed. A graphic summary of the contents of the review is presented in Figure 1. An important limitation in literature is the lack of consistent nomenclature for BEVs produced by GNB as at least two different types have been described: OIMVs, and OMVs. In this review, we termed OMVs both types, as almost all analyzed articles named them as such, and did not state which type of BEV was being discussed. This is the fundamental source of confusion in the understanding of important mechanisms for OMVs cargo loading. Additionally, many aspects of OMVs biology are still poorly described: biogenesis pathways, signaling for release, receptors in the target cell, and mechanism for host cells to respond. This hinders concluding whether or not a specific OMV cargo has meaningful biological value to explore or is the result of contamination. As a result, contradictory results between different working groups exist, and several authors have clear positions on particular aspects regarding OMVs. This article is not intended to be a critical review, but rather to present the information as it is stated, and to be an invitation for the curious and informed reader to draw their own conclusions.

Nonetheless, literature describes that OMVs derived from pathogenic bacteria, like *P. gingivalis*, contain virulence factors and have inflammatory and immunomodulatory properties. Those shed by commensal bacteria, such as OMVs from *B. fragilis* or *A. muciniphila*, play mostly an immunomodulatory role.

However, knowledge available for different species is different. Some genera have very limited information (e.g. *Odoribacter* sp.) while other dominates literature (*P. gingivalis*). Given the heterogeneity of OMVs, the characteristics of every species’ OMVs should be studied independently; therefore, there is still a broad field of research to be developed. Considering that several of the included studies were published in recent years, this subject is in constant development, and it would be worth updating the knowledge synthesis often.

The understanding of OMVs, their nature and their interactions in a biosystem not only deepens our comprehension of the bacterial reality, but also clears the path to their biotechnological use. The development of the field has the potential for vaccine production, adjuvant platforms, antibacterial treatment, biofilm studies, drug delivery, and more. Further studies are necessary to assure safe use of OMVs in the prevention and treatment of infection and disease. Finally, the study of OMVs produced by anaerobic GNB is also important for broadening our comprehension of mucosal immunity as they are produced by commensal and pathogenic bacteria.

## Funding details

P.C.-V. and L.A-A. were supported by Vicerrectoría de Investigación from Universidad de Costa Rica (VI-UCR) under grant 803-B9-455. VI-UCR had no role in study design, collection, analysis, interpretation of data, nor in writing the manuscript.

## Disclosure statement

The authors declare that they have no competing interests.

## Author contributions

**P.C.-V.:** Formal analysis, Data Curation, Writing - Original Draft, Visualization; **F.B.H.:** Data Curation, Writing - Review & Editing, Supervision; **L.A.-A.:** Conceptualization, Methodology, Validation, Writing - Review & Editing, Supervision.

## Data availability

Data will be made available on request.

## Figures, Tables and Supplementary data

**Table S1.** Summary sheet used to extract and organize the information from the articles included in the review.

| Article n° |
| --- |
| **Authors:** |
| **Publication year:** |
| **Title:** |
| **DOI:** |
| **Main objective:** |
| **Obtained information:**<br>1. Definition:<br>2. Production signals:<br>3. Biogenesis:<br>4. Content:<br>5. Exportation mechanisms:<br>6. Intracellular traffic:<br>7. Functions:<br>8. Study methods:<br>9. Other information: |
| **Pending issues:** |

## Notes

### Competing Interest Statement

The authors have declared no competing interest.

